# Altricial brains and the evolution of infant vocal learning

**DOI:** 10.1101/2024.10.29.620895

**Authors:** Renata B. Biazzi, Daniel Y. Takahashi, Asif A. Ghazanfar

## Abstract

Human infant vocal development is strongly influenced by interactions with caregivers who reinforce more speech-like sounds. This trajectory of vocal development in humans is radically different from those of our close phylogenetic relatives, Old World monkeys and apes. In these primates most closely related to humans on the evolutionary tree, social feedback plays no significant role in their vocal development. Oddly, infant marmoset monkeys, a more distantly related New World primate, do exhibit socially guided vocal learning. To explore what developmental mechanism could have evolved to account for these behavioral differences, we hypothesized that the evolution of human and marmoset vocal learning in early infancy in both species is because they are born neurally altricial relative to other primate and in a cooperative breeding social environment. Our analysis found that, indeed, human and marmoset brain are growing faster at birth when compared with chimpanzees and rhesus macaques, making them altricial relative to these primates. We formalized our hypothesis using a logistic growth model showing that the maturation of a system dependent on the rate of brain growth and the amount of social stimuli benefits from an altricial brain and a cooperative breeding environment. Our data suggest that in primates, the evolution of socially guided vocal learning during early infancy in humans and marmosets was afforded by infants with a relatively altricial brain and behavior, sustained and stimulated by cooperative breeding environments.

**Significance statement:** Humans rely on social feedback from caregivers to learn how to produce species-typical sounds, whereas other primates like macaque monkeys or chimpanzees do not. What accounts for this difference in developmental strategies? We tested the hypothesis that being born with a more immature (thus more plastic) brain may be the reason by using marmoset monkeys. This species is more distantly related to humans but exhibit the same type of vocal learning and who have a similar socially rich infant care environment. We found that, indeed, human and marmoset brain are growing faster at birth when compared with chimpanzees and rhesus macaques, making them altricial relative to these primates and this explains their similar vocal developmental strategies.

## Introduction

The prelinguistic vocal development of human infants is strongly influenced by interactions with caregivers and others. In fact, the acoustic structure of their first cries has already been influenced by the ambient prenatal linguistic environment (1). Postnatally, these cries then later differentiate into different forms which may indicate different behavioral states, soliciting varying responses by caregivers. Later, when infants begin to babble—producing consonant-vowel combinations—these, too, call the attention of caregivers (2) who may (without realizing it) reinforce more speech-like sounds than others (3). This will change the infant’s vocal output to converge more frequently upon more speech-like utterances, creating a positive feedback loop of vocal learning (4, 5). This socially guided vocal learning during infant development is an important component of the human developmental system. Yet, the mechanisms by which this early experience-dependent vocal learning occurs, and how it evolved, are still unclear.

This trajectory of vocal development in humans is radically different from those of our close phylogenetic relatives, Old World monkeys and apes. In these primates, social feedback plays no significant role in their vocal development (6, 7) (Figure 1A). So, if our closest living relatives don’t share anything like our vocal learning capacity, how did we evolve one? Humans are born altricial relative to these other primates when it comes to motor development, and a greater proportion of brain growth occurs after birth when compared with other primates (8-10). One straightforward possibility, therefore, is that humans are born with a more plastic brain than other Old World primates and this makes our earliest behaviors—like vocal production—more susceptible to the influence of others. The peculiar combination of a rapidly growing postnatal brain, a signature of altriciality (11), and the rich social environment required to care for a helpless infant may have an outsized impact on behavioral learning. This basic idea has been around for a long time (e.g., (12)) and continues to be advocated in the contemporary scientific literature (10, 13-15). For vocal learning in particular, altriciality is cited as a key factor (16, 17). Charvet and Striedter make the connection explicit for songbirds, at least (16): Altriciality in songbirds (relative to ducks and geese, for instance) allows telencephalic maturation to occur *ex ovo* which, they argue, fosters neural adaptations that help vocal learning. They suggest that this was a prerequisite for the evolution of neural circuits that allowed songbirds to produce learned vocalizations.

**Figure 1.**
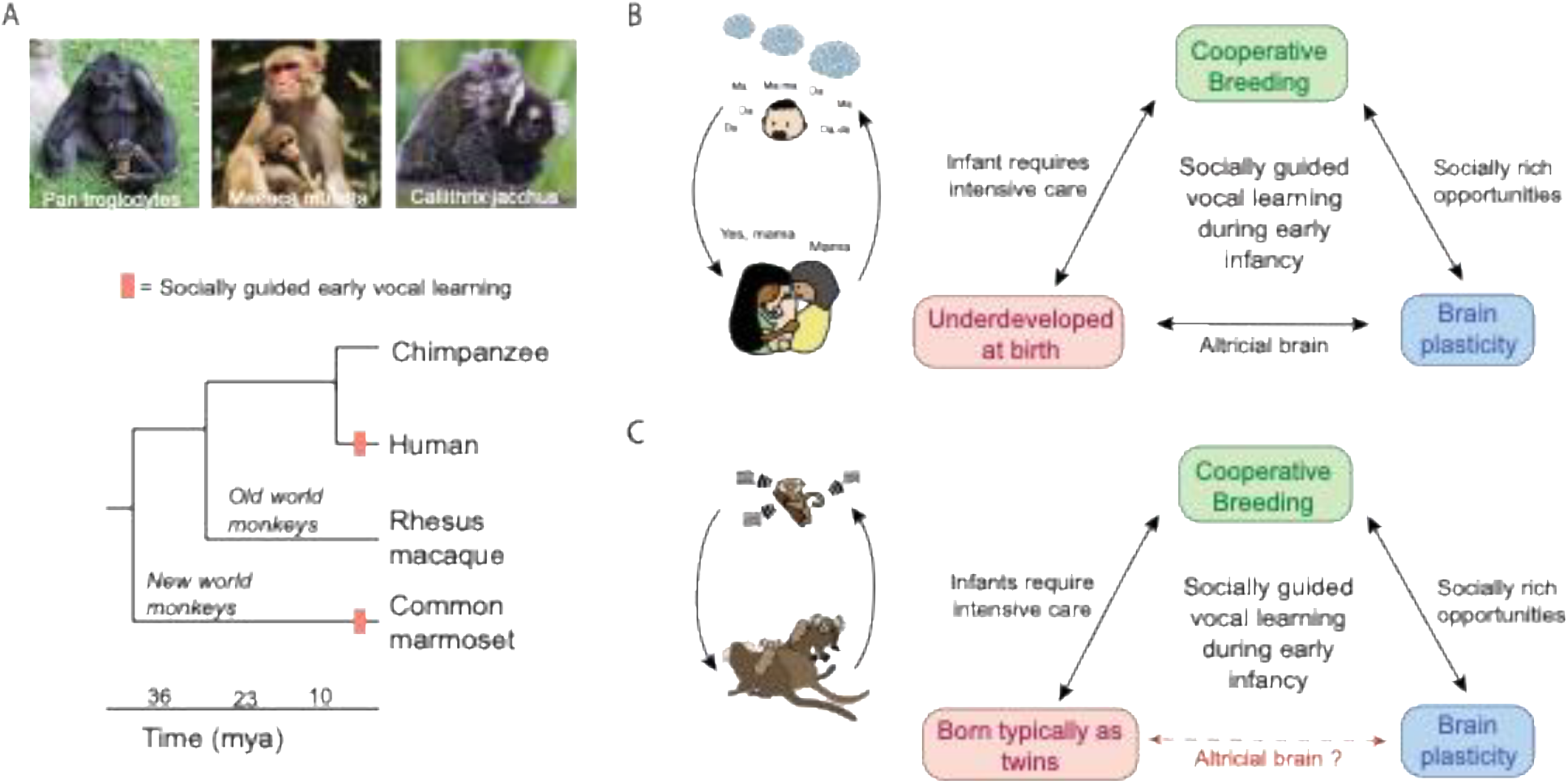
Socially guided early vocal learning hypothesis. (A) Phylogenetic tree of the four primates being compared. (B) Hypothesis about the interacting factors allowing the evolution of socially guided vocal learning during early infancy in humans. (C) Hypothesis for a case of convergent evolution of the same process in marmosets. Photo credit: Flickr users Derek Keats (Pan troglodytes) (CC BY 2.0 DEED license), Tim Ellis (Macaca mulatta) (CC BY-NC 2.0 DEED license), and Wikipedia user Nortondefeis (Callithrix jacchus) (CC BY-SA 4.0 license).

Can we actually test the idea that an altricial brain is important for the evolution of vocal learning in primates? We think so and by using marmoset monkeys as a comparative species (Figure 1A). Marmoset monkeys (*Callithrix jacchus*) are New World primates. Any behaviors they share with humans (but, importantly, not with other Old World primates) would be evidence of convergent evolution. Evidence of evolutionary convergences is an opportunity to look for common mechanisms underlying behavior across species. The interpretation of such comparisons avoids the bias created by relatedness (18, 19). Two behaviors that marmosets share with humans, but not other Old World primates, are cooperative breeding (in which both parents, older siblings and unrelated conspecifics help care for infants; (20)) and vocal learning through social feedback (21-24).

In this study, we analyze the trajectory of brain development relative to birth timing in marmoset monkeys and compare them to humans and two other Old World primates, the macaque monkey (*Macaca mulatta*) and chimpanzees (*Pan troglodytes*). We hypothesize that the trajectory of marmoset monkey brain growth relative to birth timing should be more similar to humans than to the other species. This means that their brain growth should peak around birth and not before it. We then formalize the idea that an altricial brain combined with a rich social environment enhances learning using a mathematical model.

## Results

One hypothesis is that human infants can learn to vocally communicate due to their flexible brains developing in rich social environments that drive and reinforce learning (Figure 1B) (10, 15). Their brain plasticity and susceptibility to external influences result from their altricial state at birth (13, 14). This under-development (relative to other Old World primates) makes infants highly dependent on intensive care for an extended period, a need that caregivers can supply only through social cooperation. We extend this hypothesis to marmoset monkeys, who routinely produce twins that require lots of care from mothers, fathers, siblings and others (25) (Figure 1C). The fact that marmosets are typically born as twins limits the amount of maturation that can occur inside the uterus. Consequently, they are born relying on others to carry them, and this need is met through a cooperative breeding structure involving multiple caregivers (20).

Human and monkeys share multiple motor systems for vocal production. This system must activate and coordinate laryngeal tension, respiration and articulation (26) and utilizes varying levels of cognitive control (27-29). Numerous brain regions, both subcortical and cortical, are recruited and coordinated to guide and generate vocal behavior (30-32). In light of this, we used a whole brain measure of growth to test the hypothesis that the relative altriciality of the brain at birth is important for socially guided vocal learning. We first did a literature survey to gather brain growth data from conception to the end of the juvenile stage for humans, chimpanzees, rhesus macaques, and marmoset monkeys. Figure 2A presents the data on brain volume normalized to the average adult brain size for each species. Using the combined datasets, we employed cubic splines to interpolate the available data points and estimate the brain growth curve for each species.

**Figure 2.**
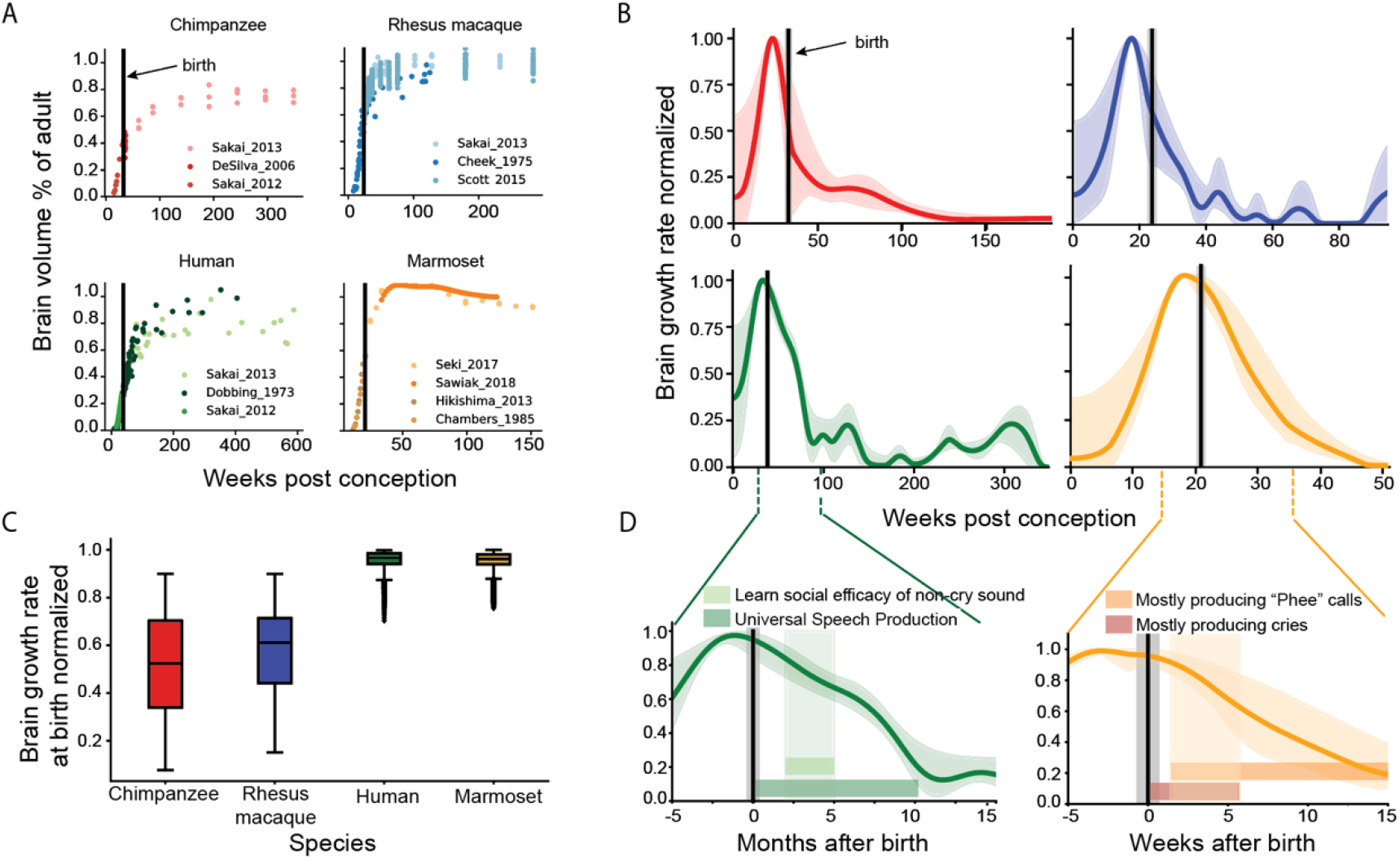
The human and marmoset brain are relatively altricial at birth compared to chimpanzees and rhesus macaques. (A) Brain volume size relative to the average adult brain size from conception to juvenile for chimpanzees, rhesus macaque, humans, and marmosets. Different shades of color indicate the source of the data. Vertical bars indicate the average birth timing for the species. (B) Brain growth rate normalized for chimpanzee (red), rhesus macaque (blue), human (green), and marmoset (orange). Vertical bars represent expected birth. Darker lines represent the average of the different fits and shaded areas contain the 2.5 and 97.5 percentile of each point in time. (C) Distributions of possible brain growth rates at birth given the birth window and the different fits for the data. Human (median = 0.96) and marmoset (median = 0.95) rates of brain growth at birth are well discriminated relative to the chimpanzee (median = 0.52) and rhesus macaque (median = 0.58) growth rates (). (D) Human (green) and marmoset (orange) early infancy vocal milestones occurring when their brains are still in a phase of rapid brain growth.

Given that different species exhibit varying growth behaviors throughout development, we choose to interpolate the data over imposing specific models that might not accurately capture each species’ unique growth trajectory. We conducted a sensitivity analysis by applying a range of parameter values in each species’ interpolation (Table 1). This approach allowed us to generate multiple potential brain growth curves for each species, reducing the risk of non-robust results due to overfitting the limited sample size. Then, by taking the curves’ derivative and filtering out obvious overfitting cases, we obtained the brain growth rate for each species (Figure 2B). Darker lines represent the average of the different fits with shaded areas containing values between the 2.5 and 97.5 percentile of each point in time. The total brain growth rate is a broad measure of the amount of change the brain is experiencing over time (changes could include neurogenesis, synaptogenesis, myelination and/or vascularization) (8, 33, 34).

**Table 1.** Parameters values for the sensitivity analysis.

| Parameters | values |
| --- | --- |
| $\lambda$ | [0.5,0.4,0.3,0.2,0.1,0.05,0.01,0.005,0.001] |
| $\omega_i$ with $i = 1,2,3$ | [1,0.75,0.5,0.25,0.1,0.05] |
| $\tau$ | [6,4,2,1,0, -1, -2, -4, -6 ] |

To analyze brain growth rate timing relative to birth, we plotted the distributions of brain growth rates at birth (Fig. 2C). We observe that the human and marmoset rates of brain growth at birth (human median = 0.96, marmoset median = 0.95) are significantly higher–-closer to the peak– - relative to the chimpanzee (median = 0.52) and rhesus macaque (median = 0.58) growth rates. We used ROC curves to test how well the distributions of each pair of species can be distinguished based on different thresholds (Fig 4C). ROC curves are plots of the proportion of “correctly predicted altricial species/all true altricial” – named True Positive Rate (TPR) – on the y-axis against the proportion of “precocial predicted as altricial/all true precocial” – named False Positive Rate (FPR) – on the x-axis. We found that marmosets and humans can be reliably differentiated from rhesus macaques, with AUC values of 0.9989 (95%CI [0.9988,0.9989]) and 0.9979 (95%CI [0.9977,0.9981]), respectively. Similarly, both species can be reliably distinguished from chimpanzees, with AUC values of 0.9985 (95%CI [0.9984,0.9985]) for marmosets and 0.9973 (95%CI [0.9972,0.9975]) for humans. However, when comparing the distributions of marmosets with humans and chimpanzees with rhesus macaque, we can’t reliably distinguish them, as they have AUC values of 0.5050 (95%CI [0.5011,0.5090]) and 0.5920 (95%CI [0.5897,0.5943]), respectively.

Using a threshold of 0.9, humans and marmosets can be differentiated from both chimpanzees and rhesus macaques with an FPR = 0 and TPR around 0.92. These finding indicates that both humans and marmoset monkeys estimates of brain growth rates at birth are similar to each other but differ from those of chimpanzees and rhesus macaques. They are still in their rapidest phase of brain growth just after birth and, relative to macaque monkeys and chimpanzees, they are neurally altricial.

**Table 2.** Cut values for brain growth fits.

| species | R2 | Non-monotonicity |
| --- | --- | --- |
| Chimpanzee | > 0.95 | > -0.5 |
| Rhesus macaque | > 0.93 | > -0.5 |
| Human | > 0.95 | > -0.5 |
| Marmoset | > 0.98 | > -0.5 |

The relatively rapid brain growth of humans and marmosets occurs in a period in which infants are experiencing important milestones of their early vocal learning (Figure 2D). For humans (green plot), some important prelinguistic milestones occurring in this period are the infant learning of the social efficacy of their non-cry sounds (2) and a phase of universal speech production just before cultural specialization (35). For marmosets (orange plot), infants experience a transition between mostly producing cries to mostly producing “phee” calls–-an important vocalization for vocal exchanges which help to maintain social contact (36), label individuals (37) and interrogate the social environment (38). Overall, these results support the hypothesis that socially guided early vocal learning milestones of humans and marmosets happen during a phase in which their brains are still rapidly developing–a period associated with increased brain plasticity (17).

Our results show that both marmosets and humans reach early vocal production milestones when their brains are in a phase of rapid brain growth, a characteristic of altricial brains (Figure 2D). From this, we hypothesize that socially guided vocal production learning during early infancy is driven by two key elements: 1) During maturation, altricial brains are more susceptible to, and dependent on, external stimuli compared to precocial brains. They are more plastic. 2) The cooperative breeding strategy required to care for helpless human infants or marmoset twins results in a very rich social environment, providing more opportunities for social learning compared to environments with primarily maternal care. To formalize our hypothesis, we developed a mathematical model designed to illustrate how varying trajectories of brain plasticity and social stimuli can influence vocal maturation through social learning (Fig. 3A). We selected a logistic differential equation for its simplicity in describing bounded growth—from 0, representing an immature stage, to 1, representing a mature stage—driven by a rate defined as (*r* x *s*). The logistic model is also one of the most standard ways to model bounded growth (39). In this model, the *r* parameter represents brain plasticity, while the *s* parameter reflects the level of social stimulus.

**Figure 3.**
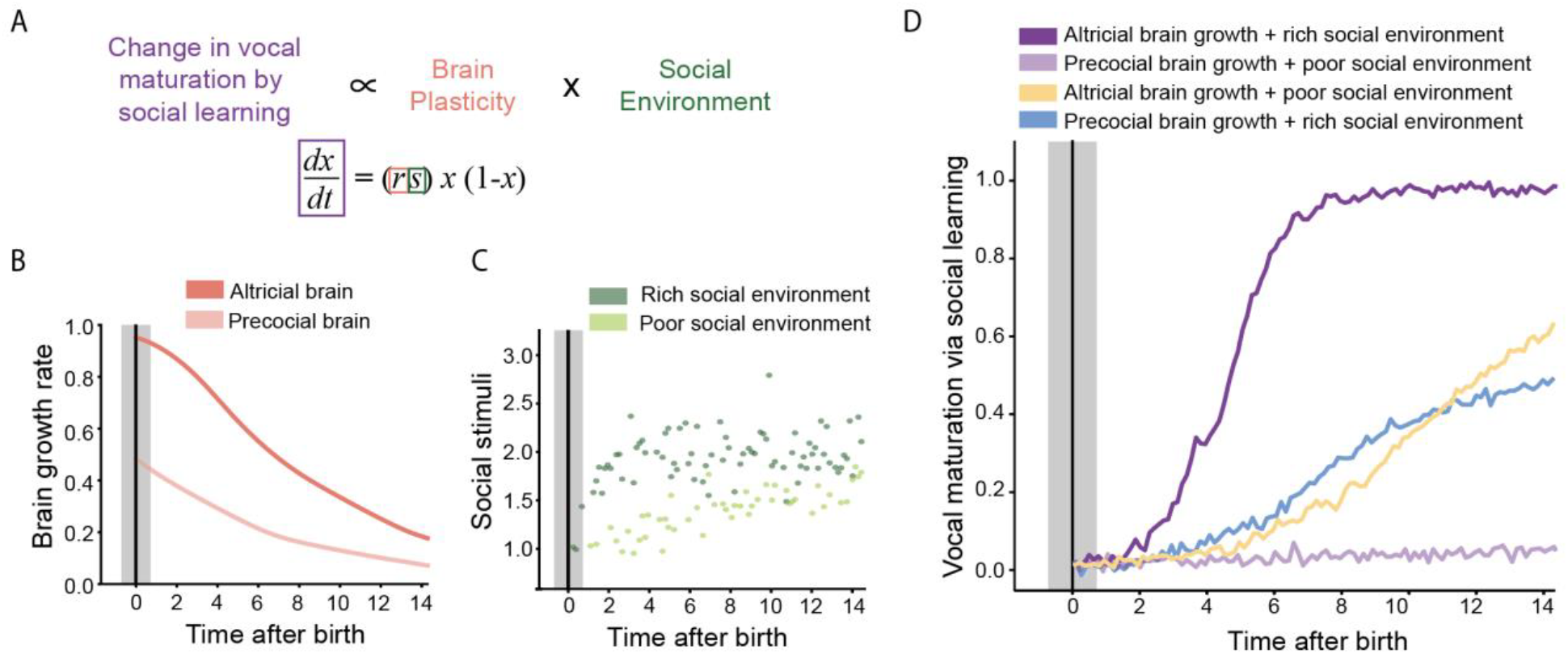
A model formalizing our hypothesis results in a system with different behavioral maturation depending on the brain growth rate and social stimuli provided during early development. (A) Formalization of the hypothesis by a logistic differential equation. The model represents a change in behavioral maturation as a function of brain plasticity and the social environment. (B) Two different brain growth rates were used to represent brain plasticity (r in the equation of Figure 3A). The altricial brain (dark pink) represents a brain growth rate like the ones of marmosets and humans. The precocial brain (light pink) represents a brain growth rate like the ones of rhesus macaques and chimpanzees. (C) The two distributions of social stimuli in time created for the simulation of vocal maturation via learning (s in the equation of Figure 3A). The dark green distribution represents an environment with frequent and intense social stimuli since early life, such as in a cooperative breeding group. The light green distribution represents an environment with less frequent and intense social stimuli early in life with the intensity increasing in time, such as infants with only maternal care. (D) Simulation of the vocal maturation using the logistic differential equation of Figure 3A with r and s values taken from Figures 3B and 3C. A system with an altricial brain and cooperative breeding (dark purple) would allow a rapid transition from immature to mature vocal production based on social learning. A system with a precocial brain and a low amount of social stimuli (light purple), on the other hand, does not allow for vocal production learning due to social stimuli. The intermediate cases – altricial brain and poor social environment (yellow) and precocial brain and rich social environment (blue) – allow intermediate levels of maturation through learning.

**Figure 4.**
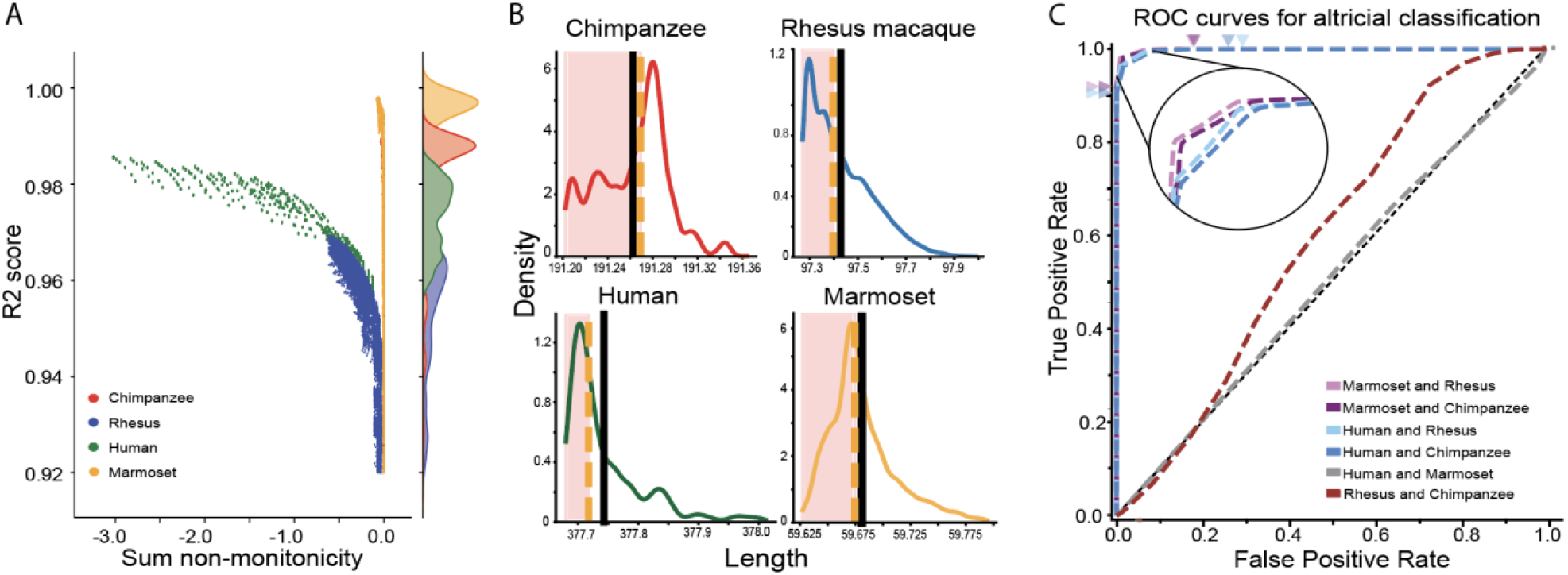
Procedures of the data analysis. (A) For each fit and each species, we calculated the R2 score for and non-monotonicity values (a measure of overfitting). Based on this plot, we defined the cut values for each species (Table 2). (B) Distribution of the lengths of each derivative curve. Black line represents the mean and the yellow dashed line represents the median. We selected only the first 50th percentile of lengths to reduce overfitting. (C) ROC curves showing how well the possible values of brain growth rate at birth can be used to find reliable thresholds to differentiate species as altricial and precocial. In each pairwise comparison, the first species is considered as the altricial one (positive classification for altriciality) and the second the precocial one (negative classification for altriciality). The arrows in the top represent the threshold in which the TPR = 1 and the FPR is minimum (threshold = 0.75 for marmosets and threshold = 0.70 for humans). The arrows in the left represent the threshold in which FPR = 0 and the TPR is maximal (threshold = 0.9 for both marmosets and humans). We see that the marmoset and human distribution is reliable to differentiate them from rhesus and chimpanzee. However, humans cannot be reliable differentiated from marmoset, neither rhesus from chimpanzees.

To represent brain plasticity, we assume that brains experiencing higher rates of growth are more plastic. Therefore, based on our fits (Figure 2B) we used an altricial (dark pink) and a precocial (light pink) regime of brain growth rates as inputs to the *r* variable in the model at each point in time (Figure 3B). To represent the social environment, we created two distributions to illustrate the amount of social experience infants have with different individuals depending on their species’ typical mode of care. Based on data from infant marmoset social experiences (40, 41), we developed a distribution depicting an early life environment containing expression of social signals from conspecifics with high frequency and consistent with a cooperative breeding environment (Figure 3C). Based on data from interactions between chimpanzee and rhesus macaque infants with other individuals in their social group (42-44), we developed a second distribution of less frequent social signals that gradually increase as the individual matures (light green). This distribution represents a less rich social environment during early infancy, reflecting the experience of infants receiving only maternal care in early infancy and slowly becoming independents and starting to interact with other conspecifics (Figure 3C).

When we simulate the scenario of an altricial brain in a rich social environment, we observe a rapid transition from immature to mature behaviors due to social learning (dark purple in Figure 3D). This non-linear transition behavior is similar to what we observe in marmoset infants’ transition between producing immature versus mature contact vocalizations (23). In the case of a less intense social environment and a precocial brain, there is no significant change in vocal behavior due to social learning, which is consistent with the lack of infant vocal production learning in rhesus macaque and chimpanzees (light purple in Figure 3D). We can interpret this model as weighting the influence of the social environment in vocal production learning via the plasticity of the brain. In the chimpanzee case, for example, intense social stimuli later in life does not allow vocal production learning if the window of plasticity for it has already closed. The intermediate combinations— altricial brains with poor social environments and precocial brains with rich social environments— (yellow and blue in Figure 3D) permit intermediate levels of maturation through learning. This occurs because we used intermediate, non extreme, values for brain plasticity and social environment as inputs in the simulation. In other words, precocial brains retain some plasticity, enabling learning in rich social environments, while the poor social environment still provides sufficient stimuli for altricial brains to learn. These gradients probably exist in different species.

## Discussion

The order of neurodevelopmental events is relatively conserved in mammals, but their duration is a fundamental source of species variation (45, 46). Shifts between birth timing and the neurodevelopmental stage, a form of heterochrony (47), can provide different windows of opportunity for environmental and social factors to affect neurodevelopment leading to different behavioral outcomes (9-11, 15, 48). We hypothesize that the evolution of human and marmoset vocal learning in early infancy was supported by their altricial brains in a cooperative breeding social environment. We found that human and marmoset brain growth rates are faster (closer to the peak) at birth when compared with chimpanzees and rhesus macaques, making them altricial relative to these primates. We formalized our hypothesis using a logistic growth model showing that the maturation of a system dependent on the rate of brain growth and the amount of social stimuli benefits from an altricial brain and a cooperative breeding environment. Importantly, we are not saying vocal learning is the only behavior affected by variations in birth timing relative to brain development—vocal learning, its presence or absence, is simply the behavior we sought to explain in this instance.

We used birth timing relative to brain growth peak to differentiate our four primate species as relatively precocial or altricial (10). Although brain proportion at birth is typically used as the criterion of altricial versus precocial, this measure has recently been pointed out as an inaccurate for comparing neurodevelopmental maturity in primates (49). We did various estimations of brain growth rate at birth by using different fitting parameter values. The distribution of possible values for rhesus macaque and chimpanzees is wider (see Figure 2C) as their birth occurs in a region with sharper decay of brain growth velocity. Even considering this higher variance, all the estimated values for these two species were lower than 0.9, while marmoset and human distributions are concentrated around 0.95. The robustness of this difference given different parameter choices made us confident that our result is not an artifact of the fitting. Future work measuring longitudinal brain growth of the same individuals for different primate species, from conception to at least late infancy, will provide even better estimates of their relative altriciality.

In light of our analysis, humans and marmoset monkeys are born with altricial brains relative to macaque monkeys and chimpanzees. Humans and marmoset monkeys both exhibit socially guided vocal learning during early infancy. Humans and marmoset monkeys are both cooperative breeders. How did these parallels between two distantly related species (again, relative to Old World primates) come to pass (50)? Let’s look at humans first. A putative evolutionary scenario for early vocal learning, and its adopted function, is the following (and note, these steps do not presume a direction of causality): 1) human infants are born altricial due to biomechanical constraints of the birth canal and energetics (51); 2) this selects for a cooperative breeding strategy to ameliorate those energetic demands (20); 3) this strategy requires that infants must elicit attention from any number of caregivers (52); and 4) the altricial, and thus more plastic, brain of the infant allows for vocal production to be influenced by the environment (i.e., through social reinforcement) which enhances the elicitation of care (15).

Now, let’s take a look at marmoset monkeys. The marmoset monkey is a small (300-400 grams, on average), New World species that is native to northeastern Brazil. They live in social groups of ∼5 to 15 individuals many of whom are related to each other. They are on a different branch of the primate evolutionary tree than humans. The marmoset monkey branch separated from the branch leading to humans about 40 million years ago, whereas the branch for Old World monkeys separated from humans 25 million years ago (53, 54). As a result, any traits shared between marmosets and humans, but not with other Old World primates, are the product of convergent evolution. One such trait is a cooperative breeding strategy. Humans and callitrichines (the taxonomic subfamily of New World monkeys that includes marmosets) are the only primates to exhibit a cooperative breeding strategy in which both parents, older siblings and unrelated conspecifics help care for infants (20). For marmoset monkeys and other callitrichines, the strategy is necessary because they are obligate producers of dizygotic twins (25). Carrying twins is, of course, more energetic costly than carrying singletons during pregnancy. Moreover, pregnant marmosets, like pregnant humans, often experience dystocia during deliver—the infants get stuck in the birth canal (55). This suggests that marmoset mothers are also operating at their energetic and physiological limits, perhaps pushing birth timing to be earlier than in other monkey or ape species. Postnatally, caring for twins is also energetically costly (e.g., feeding, carrying, etc.), exceeding what mothers can provide alone (25, 56-59). Thus, all group members help by carrying and feeding offspring (60). The multiple possible caregivers put a premium on attracting attention, as in humans. One way to do that is to produce adult-like vocal sounds. In both humans and marmosets, doing so elicits a higher probability of caregiver responses than producing immature-sounding vocalizations (61). The altricial, more plastic brain of marmoset infants facilitates the maturation, via learning, of vocalizations that sound adult-like.

Beyond (or in addition to) physical and energetic constraints, another reason why marmoset brains maybe altricial is that they evolved an odd developmental strategy as a consequence of twinning and being small: delayed timing of organogenesis during gestation. The twin embryos stop growing for a time interval such that they lag behind the development of other nonhuman primates by about three weeks (62, 63). This delay occurs during gastrulation (Carnegie Stage VII), before the closure of the neural tube between Carnegie States IX and X (64, 65). The result is altricial offspring relative to other nonhuman primates. At birth, this is reflected in their poor locomotor skills (66-70). In fact, infant marmosets are carried all the time by different group members during the first couple of weeks of postnatal life (71). Note that this delayed timing of embryogenesis in marmoset monkeys is not the same as the “embryonic diapause” seen in many other mammals. In those cases, the reversible cessation of embryo development allows animals to prolong gestation in order to give birth to offspring at a time that would increase their chances of survival. The diapause is induced either by physiological conditions in the pregnant mother or by external, seasonal effects. As far as we know, these conditions do not apply to the delayed organogenesis of marmoset monkeys.

A major (but we feel, entirely reasonable) assumption we made is that socially guided vocal learning is some function of the learning potential of the system: the plasticity of a rapidly growing brain and the amount of social stimulation received. The brain data we presented and the life history strategies of humans and marmoset monkeys are consistent with this idea. However, it could be argued that this may be one of multiple stories that could be weaved with these facts. Thus, in order to support its plausibility, we offered a formalization of the interactions between altricial brain development and a socially rich environment (Figure 3A). We used a model with a limited upper value (bounded) as we expect vocal behavior to mature and reach a limited stage equivalent to the adult average capacity (vocal behavior does not develop indefinitely). The model integrates the effects of social stimuli and brain plasticity in the learning potential by a interaction given by their simple product. As we don’t have the necessary data to test the exact mechanisms of how the interplay between social feedback and brain development works, we didn’t want to speculate on more complex interactions between these factors. The model receives a brain growth rate input (Figure 3B) based on actual brain data (Figure 2) that we defined as a proxy for the brain plasticity. It also receives a distribution of social stimuli over time (Figure 3C) that was artificially generated by distributions with varying intensities and frequencies representing cooperative and independent (e.g., female-only) breeding environments. When we simulate the scenario of an altricial brain in a rich social environment, we observe a rapid transition from immature to mature behaviors due to social learning.

Our data suggest that in primates, the evolution of socially guided vocal learning during early infancy in humans and marmosets was afforded by infants with a relatively altricial brain and behavior, sustained and stimulated by cooperative breeding environments. More data on vocal development, brain development, and breeding structure of other primates (especially New World Monkeys) is necessary to further test this hypothesis. Future research, potentially also in other classes such as birds (16), will be able to elucidate if this intertwining (in the various degrees and forms it can take in different species) is a principle behind the evolution of socially guided vocal learning during early development.

## Methods

### 1. Data collection and treatment

All the brain data for this study were collected from previously published papers, after a systematic literature review. We couldn’t find brain volume data from the entire period (from conception to late infancy) for other New World Monkeys beyond the common marmoset.

For humans, data were compiled from the total brain weight of 139 normal human brains from the 10th week post conception to the 7 postnatal years (72), the volume’s median of 203 normal human fetuses from the 18th to the 32nd week post conception (73), and total brain volume of 28 children (14 males and 14 females) from one month to 10.5 years (74).

For chimpanzees, data were compiled from the total brain volume of three growing individuals from 6 months to 6 years (74), the total brain volume of two growing fetuses from the 14th to the 33rd week post conception (73), and the neonatal brain mass of 22 newborn chimpanzees with normal body size (75).

For rhesus macaques, data were compiled from the total brain volume of six subjects (4 males and 2 females) with normal development from 3 months to 4 years (74), the total brain weight from 102 subjects from the 7th week post conception to 2 years of age (76), and the total brain volume of 44 subjects from 1 to 260 weeks of postnatal age (49).

For marmosets, data were compiled from the total brain weight of 48 subjects from 80 to 140 days of gestation (77), the total brain volume of 24 embryonic and fetal marmosets from 8 to 19 weeks post conception (78), the total brain volume of 23 subjects between 1 and 30 months (79), and the total brain volume of 41 subjects between the 3rd and 27th postnatal months (80).

All the data was collected from tables or figures available at the cited sources. In the case a table was not available, we used computer vision-assisted software to extract numerical data from plots (81). In the cases where only the brain weight was available, we converted it to the total brain volume using the average brain density (“Composition of BRAIN (ICRP),” n.d.). Subsequently, all ages were standardized to weeks post-conception using the average gestational period of each species – available at (46) for humans, chimpanzees, and rhesus macaque, and at (82) for marmosets. Finally, brain volume was normalized relative to adults using the average brain volume of adult individuals of each species (49, 83).

The fact that we had a limited sample and there is variability due to individual differences and due to different methods of measurement could have resulted in inconsistency in the brain trajectories predicted by different papers. However, despite being gathered from different sources, the data points for each species were sufficiently consistent with each other (Fig 1A). We also fitted the data using varying parameter values to estimate broad intervals for the fit, as explained in the next section.

### 2. Data fit and sensitivity analysis

We utilized univariate cubic splines to achieve smooth interpolation of our data points (Fig. 2A). The cubic spline function has a smooth parameter *λ*, which adjusts the data based on weights *ω* assigned to each point. Due to the temporal non-uniformity of our dataset – because most of the data sources we used were only from before or after birth – we opted to partition it into three distinct windows. This approach facilitated the use of varying weights for data points within each window (*ω*_1_, *ω*_2_, *ω*_3_) - thereby allowing for a more adaptable fitting process. The windows *w*_1_, *w*_2_, *w*_3_ were defined in the following way:

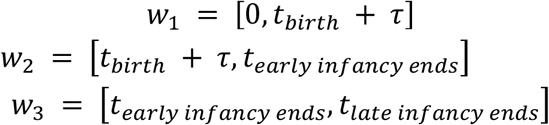

With *τ* as a parameter that varied to account for different choices of windows. This way, we performed a sensitivity analysis of our fit to the data varying the following parameters:

For every parameter combination (9 × 6^3^ × 9 = 17496), we calculated the R2 score, indicating how much variance in the data the fit explains. We also calculated the non-monotonicity, quantifying the degree of decrease in sequential points. Given that we are modeling brain growth, we expect a consistently monotonic trend (view (79) for an exception during marmoset late infancy). Therefore, we want both to maximize the variance explained (R2 score) and minimize overfitting (measured by the non-monotonicity). The non-monotonicity is the cumulative sum of the difference between decreasing consecutive values. In this phase of development, we don’t expect brain volume to decrease. Thus, any decrease in this case will be caused by the cubic-splines overfitting the data. To minimize overfit, we want the sum of non-monotonicity to be closer to zero.

Considering the cases of fits with R2 scores greater than 0.92 for all, we observed obvious cases of overfitting for humans and rhesus macaque (oscillating curves), and also that some curves for marmosets and chimpanzees were very underfitted (such as straight lines before birth). Thus, we plotted the R2 and non-monotonicity values (Figure 4A) for all the species and defined the following cut values for each one of them:

After filtering out obvious cases of overfitting and underfitting, we computed the derivatives of each curve and found evident cases of overfitting for them, as they had curves exhibiting substantial oscillations. To control for these cases, we introduced a control measure by calculating the length of each derivative curve and selecting only the first 50th percentile of lengths, as depicted in Figure 4B. This approach relies on the observation that curves with excessive oscillations, indicative of overfitting, tend to have relatively longer lengths in the distribution of curve lengths. The resulting distribution of curves for each species represents a set of estimates for brain growth derivatives that is robust to variations in the fitting parameters and were controlled for overfitting and underfitting. After the final filtering, our sample included 7.963 fits for chimpanzees, 6.764 for rhesus macaques, 2.350 for humans, and 7.005 for marmosets. We selected all the values of the rate of brain growth contained in the birth interval of each species [*t*_*birth*_ - *sd*_*birth*_, *t*_*birth*_ + *sd*_*birth*_] to generate the distributions of possible values for the rate of brain growth at birth (Fig. 2C). The size of this distribution for each species was 103545 for chimpanzees, 142086 for rhesus macaques, 16464 for humans, and 168168 for marmosets.

We generated ROC curves to evaluate how well the possible values of brain growth rate at birth can be used to find reliable thresholds to differentiate the species as altricial and precocial (Figure 4C). We compared all pairs of species: [Marmoset, Rhesus], [Marmoset, Chimpanzee], [Human, Rhesus], [Human, Chimpanzee], [Human, Marmoset], and [Rhesus, Chimpanzee]. For the comparisons, the species with the higher median – the first species in each pair – was considered the true altricial (positive classification for altriciality), while the second species was considered the true precocial species (negative classification for altriciality). To test whether the brain growth rates were distinct enough to differentiate between species, we used a binary classification using thresholds. We tested 21 thresholds with intervals of 0.05 between them: [0.00, 0.05, 0.10, …, 0.95, 1.00] (results were the same when testing 101 thresholds with intervals of 0.01). Each point of each species distribution (Fig. 2C) was predicted as altricial (positive) if its brain growth rate at birth was >= threshold, and as a precocial (negative) otherwise. Then, we created an ROC curve – which plots the proportion of “correctly predicted altricial species/all true altricial” on the y-axis against the proportion of “precocial predicted as altricial/all true precocial” on the x-axis. The arrows in the superior part of the plot represent the thresholds in which TPR = 1 and the FPR is minimum (threshold = 0.75 for marmosets and threshold = 0.70 for humans). The arrows in the left part of the plot represent the thresholds in which FPR = 0 and the TPR is maximal (threshold = 0.9 for both marmosets and humans in all their comparisons). Thus, marmosets’ and humans’ distributions are reliable to differentiate them from rhesus macaques and chimpanzees. However, humans cannot be reliable differentiated from marmoset, neither rhesus from chimpanzees.

The area under the curve (AUC) represents how well the two distributions can be differentiated. An AUC of 1 indicates that the distributions can be perfectly differentiated 100% of the time, with no overlap between them. An AUC of 0.5 implies there is a 50% chance of correctly classifying points in the two distributions, representing a classification with the same change of an unbiased random guessing. We did a bootstrapping of N = 1000 in each pairwise comparisons to estimate a confidence interval for their AUC. Marmosets and humans can be reliably differentiated from rhesus macaques, with AUC values of 0.9989 (95%CI [0.9988,0.9989]) and 0.9979 (95%CI [0.9977,0.9981]), respectively. They can also be reliably distinguished from chimpanzees, with AUC values of 0.9985 (95%CI [0.9984,0.9985]) for marmosets and 0.9973 (95%CI [0.9972,0.9975]) for humans. However, the distributions of marmosets and humans and of chimpanzees and rhesus macaque can’t be reliably distinguished, with AUC values of 0.5050 (95%CI [0.5011,0.5090]) and 0.5920 (95%CI [0.5897,0.5943]), respectively.

For Figure 2B, we applied a Gaussian smoothness for visualization purposes. The smooth parameters were set to 1 for marmosets and chimpanzees, and 10 for humans and rhesus macaques. All the code used for this analysis is available at https://github.com/biagzi/altricial_brain_vocal_learning.

## 3. Model

### a. Model definition

To formalize our hypothesis, we required a model to describe the vocal production learning depending on the plasticity of the system (*r*) interacting with social inputs (*s*). We opted to employ a logistic differential equation for this purpose, given its bounded nature and the premise that vocal learning should incrementally increases proportionally to the positive interaction between variables *r* and *s*. Consequently, in our formalization, social stimulus and plasticity (learning rate of the system) mediate one another. While other formalizations could potentially work as effective models, we opted not to impose more complex and arbitrary relationships in the variables. We also added a small Gaussian noise *ε* for the increment in learning. Thus, the differential equation defining the dynamics of x, the vocal production maturation via social learning, is:

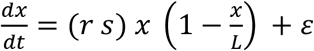

The parameter L (the carrying capacity of the system) was defined as 1 as the vocal maturation goes from 0 (immature vocalization) to 1 (mature vocalization). This equation has a direct analytic solution under the condition that r and s do not vary with time. However, in our case, we assume that the learning rate of the system corresponds to the brain growth rate, hence r = r(t), and the social stimuli fluctuate over time, denoted as s = s(t). Consequently, we solved this equation numerically.

### b. Model simulation

To numerically solve the differential equation, we used parameters based on meaningful values for the marmoset case. We adopted a time window of T = 60 (representing weeks) with a time step of dt = 1/7 (equating to one day). The simulation time starts at birth and ends after 420 time steps. The noise *ε*(*t*) was sampled from a normal distribution *N* ∼ (0,0.01). All the code used for the simulation is available at:

https://github.com/biagzi/altricial_brain_vocal_learning.

We conducted simulations for two distinct regimes of brain growth rate *r*(*t*). The first regime mirrors the altricial pattern, akin to the results found for marmosets and humans. For this simulation, we used a smoothed version of the marmoset brain growth rate average fit. In contrast, the second regime reflects the precocial pattern, resembling the results we found for the chimpanzees and rhesus macaques. For this simulation, we used the same altricial regime but temporally shifted by 7 weeks. Consequently, at birth, the rate stands at approximately 0.5, a value consistent with findings in rhesus macaques and chimpanzees.

We also established two distinct regimes for the social stimulus *s*(*t*), each aimed at emulating different social environments. The first regime mirrors an environment characterized by frequent and intense social interactions since early in life, akin to cooperative breeding observed in species like marmosets and humans. This regime was based on the percentage of time marmoset infants spend with different caregivers (40, 41).

The second regime simulates an environment with poorer social stimuli, featuring less frequency and intensity compared to the first regime. The intense of social stimuli increases in time, as we expect from infants that receive only maternal care but start to interact more with other adults as they grow old in their social groups. This regime was based on the frequency of social play (42) of infant chimpanzees and the time infant rhesus macaque mounted in other group members (43) and time spent with their mothers and peers(44).

To generate the distributions of stimuli over time for each regime, we utilized the same models with different parameters. For each time point of our simulation, we used a Bernoulli distribution with probability p of having a social stimulus and 1-p of not having. Subsequently, conditioned on the presence of a stimulus in the given time step, we sampled the stimulus intensity from a Log-Normal distribution with a mean *μ* = 0.7 and standard deviation *σ* = 0.1 for both species. To generate stimuli intensities that increase with time, we used a logistic growth curve with *k* as the growth rate. The specific parameters utilized for each regime are detailed in Table 3. For all the simulations, we used the same random seed of 1.

**Table 3.** Social stimulus parameters.

|  | Poor social environment | Rich social environment |
| --- | --- | --- |
| $p$ | 0.5 | 0.8 |
| $k$ | 0.02 | 0.2 |

## References

1. B. Mampe, A. D. Friederici, A. Christophe, K. Wermke, Newborns’ cry melody is shaped by their native language. Curr Biol 19, 1994–1997 (2009).

2. S. L. Elmlinger, J. A. Schwade, L. Vollmer, M. H. Goldstein, Learning how to learn from social feedback: The origins of early vocal development. Developmental Science 26, e13296 (2023).

3. S. L. Elmlinger, J. A. Schwade, M. H. Goldstein, The ecology of prelinguistic vocal learning: parents simplify the structure of their speech in response to babbling. Journal of Child Language 46, 998–1011 (2019).

4. M. H. Goldstein, A. P. King, M. J. West, Social interaction shapes babbling: Testing parallels between birdsong and speech. Proceedings of the National Academy of Sciences 100, 8030–8035 (2003).

5. M. H. Goldstein, J. A. Schwade, Social feedback to infants’ babbling facilitates rapid phonological learning. Psychol Sci 19, 515–523 (2008).

6. M. Belyk, S. Brown, The origins of the vocal brain in humans. Neuroscience & Biobehavioral Reviews 77, 177–193 (2017).

7. S. E. R. Egnor, M. D. Hauser, A paradox in the evolution of primate vocal learning. Trends in Neurosciences 27, 649–654 (2004).

8. J. H. Gilmore, R. C. Knickmeyer, W. Gao, Imaging structural and functional brain development in early childhood. Nature Reviews Neuroscience 19, 123–137 (2018).

9. A. Gómez-Robles, C. Nicolaou, J. B. Smaers, C. C. Sherwood, The evolution of human altriciality and brain development in comparative context. Nat Ecol Evol 8, 133–146 (2024).

10. C. C. Sherwood, A. Gómez-Robles, Brain Plasticity and Human Evolution. Annual Review of Anthropology 46, 399–419 (2017).

11. A. C. Halley, Minimal variation in eutherian brain growth rates during fetal neurogenesis. Proceedings of the Royal Society B: Biological Sciences 284, 20170219 (2017).

12. A. Portmann, A Zoologist Looks at Humankind, New York 1990. orig. Zoologie und das neue Bild vom Menschen. Hamburg (1960).

13. E. I. Knudsen, Sensitive Periods in the Development of the Brain and Behavior. Journal of Cognitive Neuroscience 16, 1412–1425 (2004).

14. B. Kolb, R. Gibb, Brain Plasticity and Behaviour in the Developing Brain. J Can Acad Child Adolesc Psychiatry 20, 265–276 (2011).

15. B. L. Finlay, R. Uchiyama, “The Timing of Brain Maturation, Early Experience, and the Human Social Niche” in Evolution of Nervous Systems. (Elsevier, 2017), pp. 123–148.

16. C. J. Charvet, G. F. Striedter, Developmental modes and developmental mechanisms can channel brain evolution. Frontiers in Neuroanatomy 5, 4 (2011).

17. J. F. Werker, T. K. Hensch, Critical Periods in Speech Perception: New Directions. Annual Review of Psychology 66, 173–196 (2015).

18. P. S. Katz, Neural mechanisms underlying the evolvability of behaviour. Philosophical Transactions of the Royal Society B: Biological Sciences 366, 2086–2099 (2011).

19. R. Powell, Contingency and Convergence: Toward a Cosmic Biology of Body and Mind (The MIT Press, 2020).

20. J. M. Burkart, S. B. Hrdy, C. P. Van Schaik, Cooperative breeding and human cognitive evolution. Evolutionary Anthropology: Issues, News, and Reviews: Issues, News, and Reviews 18, 175–186 (2009).

21. Y. B. Gultekin, S. R. Hage, Limiting parental feedback disrupts vocal development in marmoset monkeys. Nat Commun 8, 14046 (2017).

22. Y. B. Gultekin, S. R. Hage, Limiting parental interaction during vocal development affects acoustic call structure in marmoset monkeys. Science Advances 4, eaar4012 (2018).

23. D. Y. Takahashi et al., The developmental dynamics of marmoset monkey vocal production. Science 349, 734–738 (2015).

24. D. Y. Takahashi, D. A. Liao, A. A. Ghazanfar, Vocal Learning via Social Reinforcement by Infant Marmoset Monkeys. Curr Biol 27, 1844–1852.e1846 (2017).

25. R. A. Harris et al., Evolutionary genetics and implications of small size and twinning in callitrichine primates. Proc Natl Acad Sci U S A 111, 1467–1472 (2014).

26. A. A. Ghazanfar, D. Rendall, Evolution of human vocal production. Current Biology 18, R457–R460 (2008).

27. D. A. Liao, Y. S. Zhang, L. X. Cai, A. A. Ghazanfar, Internal states and extrinsic factors both determine monkey vocal production. Proceedings of the National Academy of Sciences 115, 3978–3983 (2018).

28. T. Pomberger, C. Risueno-Segovia, Y. B. Gultekin, D. Dohmen, S. R. Hage, Cognitive control of complex motor behavior in marmoset monkeys. Nat Commun 10, 3796 (2019).

29. T. Pomberger, C. Risueno-Segovia, J. Löschner, S. R. Hage, Precise motor control enables rapid flexibility in vocal behavior of marmoset monkeys. Current Biology 28, 788–794. e783 (2018).

30. S. R. Hage, A. Nieder, Dual neural network model for the evolution of speech and language. Trends in neurosciences 39, 813–829 (2016).

31. G. Holstege, H. H. Subramanian, Two different motor systems are needed to generate human speech. Journal of Comparative Neurology 524, 1558–1577 (2016).

32. Y. S. Zhang, A. A. Ghazanfar, A hierarchy of autonomous systems for vocal production. Trends in neurosciences 43, 115–126 (2020).

33. J. L. Wallace, A. A. Pollen, Human neuronal maturation comes of age: cellular mechanisms and species differences. Nature Reviews Neuroscience 25, 7–29 (2024).

34. B. D. Semple, K. Blomgren, K. Gimlin, D. M. Ferriero, L. J. Noble-Haeusslein, Brain development in rodents and humans: Identifying benchmarks of maturation and vulnerability to injury across species. Prog Neurobiol 106, 1–16 (2013).

35. P. K. Kuhl, Early language acquisition: cracking the speech code. Nature Reviews Neuroscience 5, 831–843 (2004).

36. D. Y. Takahashi, D. Z. Narayanan, A. A. Ghazanfar, Coupled Oscillator Dynamics of Vocal Turn-Taking in Monkeys. Current Biology 23, 2162–2168 (2013).

37. G. Oren et al., Vocal labeling of others by nonhuman primates. Science 385, 996–1003 (2024).

38. T. T. Varella, D. Y. Takahashi, A. A. Ghazanfar, Active sampling as an information seeking strategy in primate vocal interactions. Commun Biol 7, 1098 (2024).

39. A. Tsoularis, J. Wallace, Analysis of logistic growth models. Mathematical Biosciences 179, 21–55 (2002).

40. M. d. F. Arruda, M. E. Yamamoto, O. F. Bueno, Interactions between parents and infants, and infants-father separation in the common marmoset (Callithrix jacchus). Primates 27, 215–228 (1986).

41. J. Locke-Haydon, N. Chalmers, The development of infant-caregiver relationships in captive common marmosets (Callithrix jacchus). International Journal of Primatology 4, 63–81 (1983).

42. W. Jens, J. A. Van Hooff, R. P. Spijkerman, H. Dienske, Differences in play development of young chimpanzees reared in family groups and in peer groups. Netherlands Journal of Zoology 45, 402–421 (1994).

43. G. R. Brown, A. F. Dixson, The development of behavioural sex differences in infant rhesus macaques (Macaca mulatta). Primates 41, 63–77 (2000).

44. A. M. Ryan et al., New approaches to quantify social development in rhesus macaques (Macaca mulatta): Integrating eye tracking with traditional assessments of social behavior. Developmental Psychobiology 62, 950–962 (2020).

45. C. J. Charvet, K. Ofori, C. Falcone, B. A. Rigby Dames, Transcription, structure, and organoids translate time across the lifespan of humans and great apes. PNAS Nexus 2, pgad230 (2023).

46. A. D. Workman, C. J. Charvet, B. Clancy, R. B. Darlington, B. L. Finlay, Modeling Transformations of Neurodevelopmental Sequences across Mammalian Species. J. Neurosci. 33, 7368–7383 (2013).

47. L. R. Fenlon, Timing as a Mechanism of Development and Evolution in the Cerebral Cortex. Brain Behav Evol 97, 8–32 (2022).

48. R. Uchiyama, M. Muthukrishna, “Cultural Evolutionary Neuroscience” in Oxford Handbook of Cultural Neuroscience and Global Mental Health, J. Y. Chiao, S.-C. Li, R. Turner, S. Y. Lee-Tauler, B. A. Pringle, Eds. (Oxford University Press, 2021), pp. 0.

49. J. A. Scott et al., Longitudinal analysis of the developing rhesus monkey brain using magnetic resonance imaging: birth to adulthood. Brain Struct Funct 221, 2847–2871 (2016).

50. T. T. Varella, A. A. Ghazanfar, Cooperative care and the evolution of the prelinguistic vocal learning. Developmental Psychobiology 63, 1583–1588 (2021).

51. H. M. Dunsworth, A. G. Warrener, T. Deacon, P. T. Ellison, H. Pontzer, Metabolic hypothesis for human altriciality. Proceedings of the National Academy of Sciences 109, 15212–15216 (2012).

52. K. Zuberbühler, Cooperative breeding and the evolution of vocal flexibility. (2011).

53. T. M. Preuss, “Primate brain evolution.” in Evolutionary neuroscience J. H. Kaas, Ed. (Academic Press, Oxford, U.K., 2009), pp. 793–825.

54. T. M. Preuss, “Primate brain evolution.” in Evolutionary neuroscience., J. H. Kaas, Ed. (Academic Press, 2009), pp. 793–825.

55. L. Divincenti Jr, A. D. Miller, D. J. Knoedl, J. F. Mitchell, Uterine Rupture in a Common Marmoset (Callithrix jacchus). Comparative Medicine 66, 254–258 (2016).

56. R. A. Harris et al., Evolutionary genetics and implications of small size and twinning in callitrichine primates. Proceedings of the National Academy of Sciences 111, 1467–1472 (2014).

57. W. Leutenegger, Monogamy in callitrichids: a consequence of phyletic dwarfism? International Journal of Primatology 1, 95–98 (1980).

58. A. W. Goldizen, A comparative perspective on the evolution of tamarin and marmoset social systems. International Journal of Primatology 11, 63–83 (1990).

59. N. O. Zweifel, M. J. Z. Hartmann, Defining “active sensing” through an analysis of sensing energetics: homeoactive and alloactive sensing. Journal of Neurophysiology 124, 40–48 (2020).

60. L. J. Digby, S. F. Ferrari, W. Saltzman, The role of competition in cooperatively breeding species. Primates in perspective. Oxford University Press, New York, 85–106 (2006).

61. D. Y. Takahashi, A. R. Fenley, A. A. Ghazanfar, Early development of turn-taking with parents shapes vocal acoustics in infant marmoset monkeys. Philosophical Transactions of the Royal Society B: Biological Sciences 371, 1–12 (2016).

62. G. Marroig, J. M. Cheverud, A comparison of phenotypic variation and covariation patterns and the role of phylogeny, ecology, and ontogeny during cranial evolution of New World monkeys. Evolution 55, 2576–2600 (2001).

63. S. Montgomery, N. Mundy, Parallel episodes of phyletic dwarfism in callitrichid and cheirogaleid primates. Journal of Evolutionary Biology 26, 810–819 (2013).

64. I. R. Phillips (1976) The embryology of the common marmoset (Callithrix jacchus) - PubMed.

65. R. R. Soman et al., High resolution dynamic ultrasound atlas of embryonic and fetal development of the common marmoset. Journal of Assisted Reproduction and Genetics 41, 1319–1328 (2024).

66. M. L. Gustison, J. I. Borjon, D. Y. Takahashi, A. A. Ghazanfar, Vocal and locomotor coordination develops in association with the autonomic nervous system. eLife 8 (2019).

67. Y. Wang, Q. Fang, N. Gong, Motor assessment of developing common marmosets. Neurosci Bull 30, 387–393 (2014).

68. N. Schultz-Darken, K. M. Braun, M. E. Emborg, Neurobehavioral development of common marmoset monkeys. Developmental Psychobiology 58, 141–158 (2016).

69. M. L. Gustison, J. I. Borjon, D. Y. Takahashi, A. A. Ghazanfar, Vocal and locomotor coordination develops in association with the autonomic nervous system. eLife 8, e41853 (2019).

70. N. Schultz-Darken, K. M. Braun, M. E. Emborg, Neurobehavioral development of common marmoset monkeys. Developmental Psychobiology 58, 141–158 (2016).

71. C. T. Snowdon, Social processes in communication and cognition in callitrichid monkeys: a review. Anim Cogn 4, 247–257 (2001).

72. J. Dobbing, J. Sands, Quantitative growth and development of human brain. Archives of Disease in Childhood 48, 757–767 (1973).

73. T. Sakai et al., Fetal brain development in chimpanzees versus humans. Current Biology 22, R791–R792 (2012).

74. T. Sakai et al., Developmental patterns of chimpanzee cerebral tissues provide important clues for understanding the remarkable enlargement of the human brain. Proceedings of the Royal Society B: Biological Sciences 280, 20122398 (2013).

75. J. Desilva, J. Lesnik, Chimpanzee neonatal brain size: Implications for brain growth in Homo erectus. Journal of Human Evolution 51, 207–212 (2006).

76. D. B. Cheek, Fetal and Postnatal Cellular Growth (Wiley, London., 1975).

77. P. L. Chambers, J. P. Hearn, Embryonic, foetal and placental development in the Common marmoset monkey (<i>Callithrix jacchus</i>). Journal of Zoology 207, 545–561 (1985).

78. K. Hikishima et al., Atlas of the developing brain of the marmoset monkey constructed using magnetic resonance histology. Neuroscience 230, 102–113 (2013).

79. F. Seki et al., Developmental trajectories of macroanatomical structures in common marmoset brain. Neuroscience 364, 143–156 (2017).

80. S. J. Sawiak et al., Trajectories and Milestones of Cortical and Subcortical Development of the Marmoset Brain From Infancy to Adulthood. Cerebral Cortex 28, 4440–4453 (2018).

81. A. Rohatgi (2024) WebPlotDigitizer.

82. S. D. Tardif et al., Reproduction in captive common marmosets (Callithrix jacchus). Comparative Medicine 53, 364–368 (2003).

83. K. R. Rosenberg, The Evolution of Human Infancy: Why It Helps to Be Helpless. Annual Review of Anthropology 50, 423–440 (2021).

